# Abundance data applied to a novel model invertebrate host sheds new light on parasite community assembly in nature

**DOI:** 10.1101/2020.06.26.173542

**Authors:** Joshua I. Brian, David C. Aldridge

**Author notes:** Correspondence: Joshua I. Brian.

## Abstract

Understanding how environmental drivers influence the construction of parasite communities, in addition to how parasites may interact at an infracommunity level, are fundamental requirements for the study of parasite ecology. Knowledge of how parasite communities are assembled will help to predict the risk of parasitism for hosts, and model how parasite communities may change under variable conditions. However, studies frequently rely on presence-absence data and examine multiple host species or sites, metrics which may be too coarse to characterise nuanced within-host patterns. Here, we utilise a novel host system, the freshwater mussel *Anodonta anatina*, to investigate how both the presence and abundance of 14 parasite taxa correlate with environmental drivers across 720 replicate parasite infracommunities. Using both redundancy analysis and a joint species distribution model, we model the impact of both host-level and environment-level characteristics on parasite structure, as well as parasite-parasite correlations after accounting for all other factors. We demonstrate that both niche- and neutral-based factors are important but to varying degrees across parasite species, suggesting that applying generalities to parasite community construction is too simplistic. Further, we show that presence-absence data fails to capture important density-dependent effects of parasite load for parasites with high abundance. Finally, we highlight that predicted parasite interaction networks vary greatly depending on whether abundance or presence-absence data is used. Our results emphasise the multi-faceted nature of parasite community assembly, and that future studies require careful consideration of the data used to infer community structure.

## Introduction

Predicting the distribution and abundance of organisms is a central goal of ecology (Gravel et al. 2006). Because species do not exist in isolation, understanding how ecological communities are assembled and maintained has been and continues to be of large interest (e.g. Gleason 1927; Clements 1936; Diamond 1975; Tilman 1977; Hubbell 1997). Elucidating the factors determining community assembly is important for the management and conservation of species, and for predicting how communities may respond to disturbance such as establishment of an invasive species or large-scale environmental change (e.g. Davis et al. 2019). Individual organisms can also constitute a community of parasitic or pathogenic taxa (Pederson and Fenton 2007; Telfer et al 2010; Rynkiewicz et al. 2015). Understanding the drivers of these communities is crucial (Budischak et al 2016; Rynkiewicz et al. 2019) to enable predictions based on host and environmental characteristics (Johnson et al. 2015), thus allowing analysis of the possible effects of parasite communities on the host organism and the wider host population. Further, viewing the individual host as an ecosystem allows one to learn, with high replication, how communities interact and are shaped by environmental factors.

Assessing parasite community interactions is a challenge. Ideally, the whole suite of an organism’s parasites should be studied to accurately map community patterns and understand the effect on the host (Vaumourin et al. 2015; Fountain-Jones et al. 2019). However, until recently, only one-to-one parasite interactions have been explored (Griffiths et al. 2014; Hellard et al. 2015). When entire communities are considered, analysis frequently spans multiple host species and sites, which are often found as the greatest determinants of parasite community structure (e.g. Vidal-Martinez and Poulin 2003; Dallas and Presley 2014; Dallas et al. 2019). These effects may mask more nuanced parasite patterns and associations within hosts (a parasite ‘infracommunity’, Bush et al. 1997). As all parasite community data trends (among host species or sites) are driven by infracommunity-level interactions (Bush et al 1997; Pederson and Fenton 2007), understanding influences on infracommunity structure is key to the study of parasite community ecology. However, there is still little research with this focus (Pilosof et al. 2015; Halliday et al. 2017).

When parasite communities have been studied, investigations overwhelmingly focus on mammalian hosts, particularly rodents and ruminants (e.g. Watve and Sukumar 1995; Telfer et al. 2010; Pilosof et al. 2015; Henrichs et al. 2016; Beechler et al. 2019; Dallas et al. 2019; see review in Ezenwa 2016). However, the efficacy of these models is questionable. Mammals possess sophisticated immune systems and exhibit complex behaviour. Adaptive immune systems make it very difficult to accurately predict parasite effects (Bordes and Morand 2009; Griffiths et al. 2014; Benesh and Kalbe 2016), and can lead to cascades of immune-mediated interactions (Gorsich et al. 2018), while intraspecific behavioural diversity can lead to some individuals being disproportionately at risk, or to show ‘super-spreader’ behaviour (Paull et al. 2012; Habig et al. 2019). While these factors are important for individual species, using them as models may preclude identification of possible generalities in parasite community construction. Further, the use of mammals also generally necessitates the use of presence-absence parasite data (Hellard et al. 2015), as destructive quantitative sampling is often untenable. This means that the key tenet of ecology - to understand distribution and *abundance* – can only ever be half-addressed. However, consideration of abundance is crucial, both for parasites’ effects on hosts and also how the parasites may interact (Budischak et al. 2016; Rynkiewicz et al. 2019). Indeed, theoretical work has shown that the direction of parasite interactions may reverse based on the abundance of one of the partners (Fenton 2013), and that presence-absence data is unreliable at detecting parasite interactions (Blanchet et al. 2020).

A way forward may be an increased focus on invertebrates, which have been severely neglected in parasite community studies (Wilson et al. 2015). Invertebrates largely lack an adaptive immune system, meaning evidence for parasite infection trends and interactions can be taken at face value (Anderson and May 1981). Reduced ethical requirements for non-cephalised invertebrates facilitates abundance sampling, and invertebrates frequently harbour more macroparasites than their vertebrate counterparts (Wilson et al. 2015). For non-motile invertebrate species, the complexities of behaviour are also avoided.

This paper introduces and takes advantage of one such invertebrate system. The duck mussel *Anodonta anatina* is a sessile unionid bivalve found commonly in freshwater environments around Europe. It is infected by a broad range of parasites from multiple phyla (Brian and Aldridge 2019), and it has clear discrete tissues (e.g. mantle, gills, gonad) that are each targeted by multiple parasites, analogous to niches in typical communities. Focusing on this species at a single site, we examine factors relating to the composition of replicate infracommunities, providing a sound theoretical basis for more extensive work across larger scales. Rather than focus on a subset of parasites, our analysis incorporates both presence-absence and abundance information for all parasites observed in *A. anatina* over the course of a full year’s sampling. By modelling a range of potential environmental drivers and parasite-parasite interactions using both presence-absence and abundance frameworks, we show that different parasites respond to different factors. Further, we explicitly highlight the differences between abundance and presence-absence data, demonstrating that accounting for the abundance of parasites can lead to very different conclusions about environmental drivers and parasite interactions than if presence-absence data are used.

## Methods

### Study site and sampling

Sixty individuals of the duck mussel *Anodonta anatina* (Linnaeus 1758) were sampled from the same location at the Old West River at Stretham (52.3343° N, 0.2243°E), a lower reach of the River Great Ouse (UK), at monthly intervals from February 2019 to February 2020 (12 collections, N = 720 mussels). Mussels were sampled by hand from the river margin. This sampling size (60) was selected as it gives a >95% chance of detecting parasites present in the mussel population at 5% prevalence (Grizzle and Brunner 2009). This monthly sampling did not affect the overall density of mussels present, as the time taken to collect 60 mussels did not vary from month to month. The size distribution of mussels was also unaffected by the sampling regime (63.9 ± 11.1 mm [overall mean ± 1 s.d.], see Figure S1.1). Mussels were transported to the laboratory, and were dissected systematically to identify and count all parasites present (see S1 Supplementary Methods for full details).

### Analysis variables

Altogether, 14 parasites were included in the analyses, with two associated categorical parasite traits (Table 1): life history (whether the mussel is the only host, or whether the parasite utilises multiple hosts), and location (whether the parasite is present in the gills, mantle, or gonad of the mussel). These trait values were utilised in the Joint Species Distribution Modelling (see below).

**Table 1:**
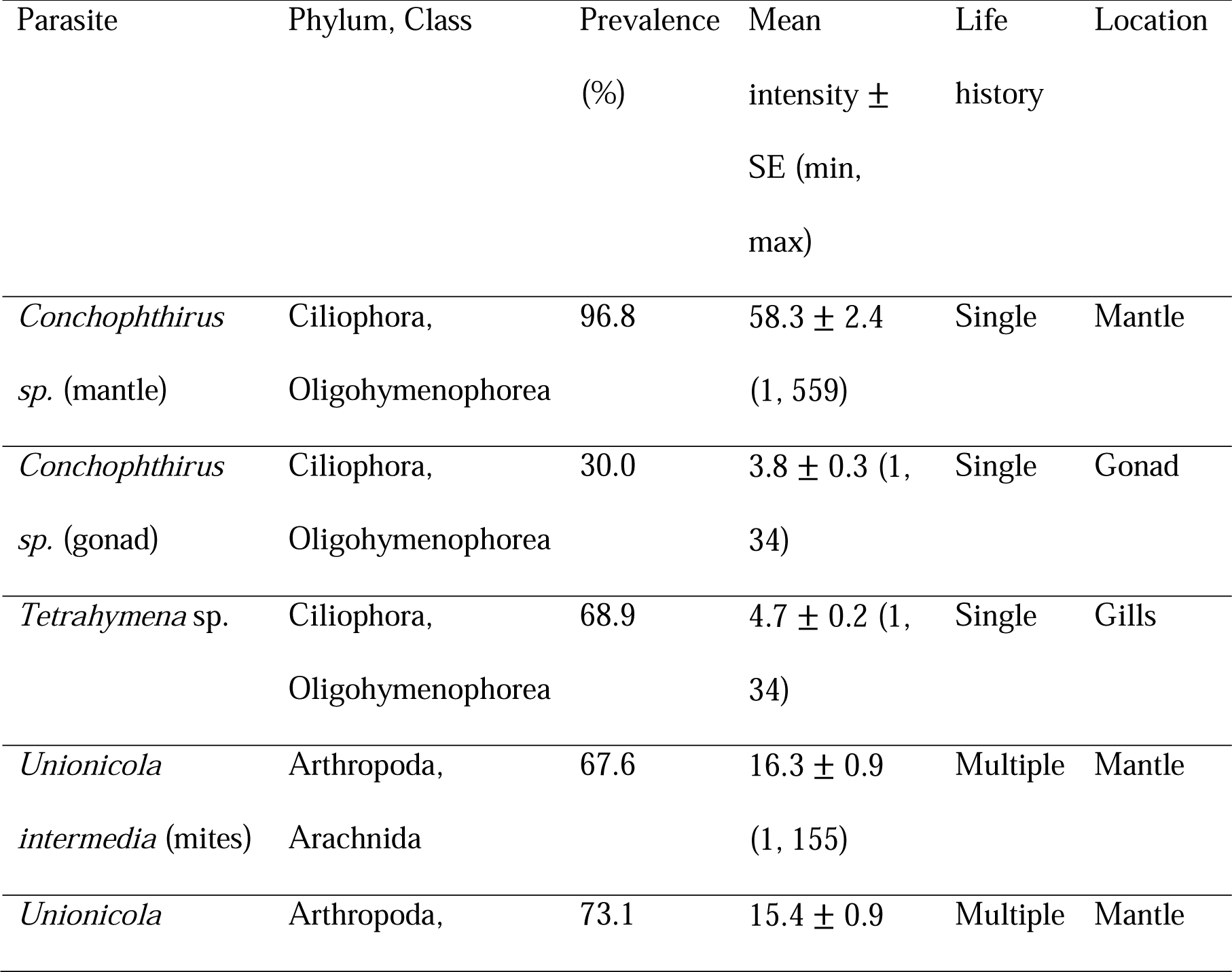
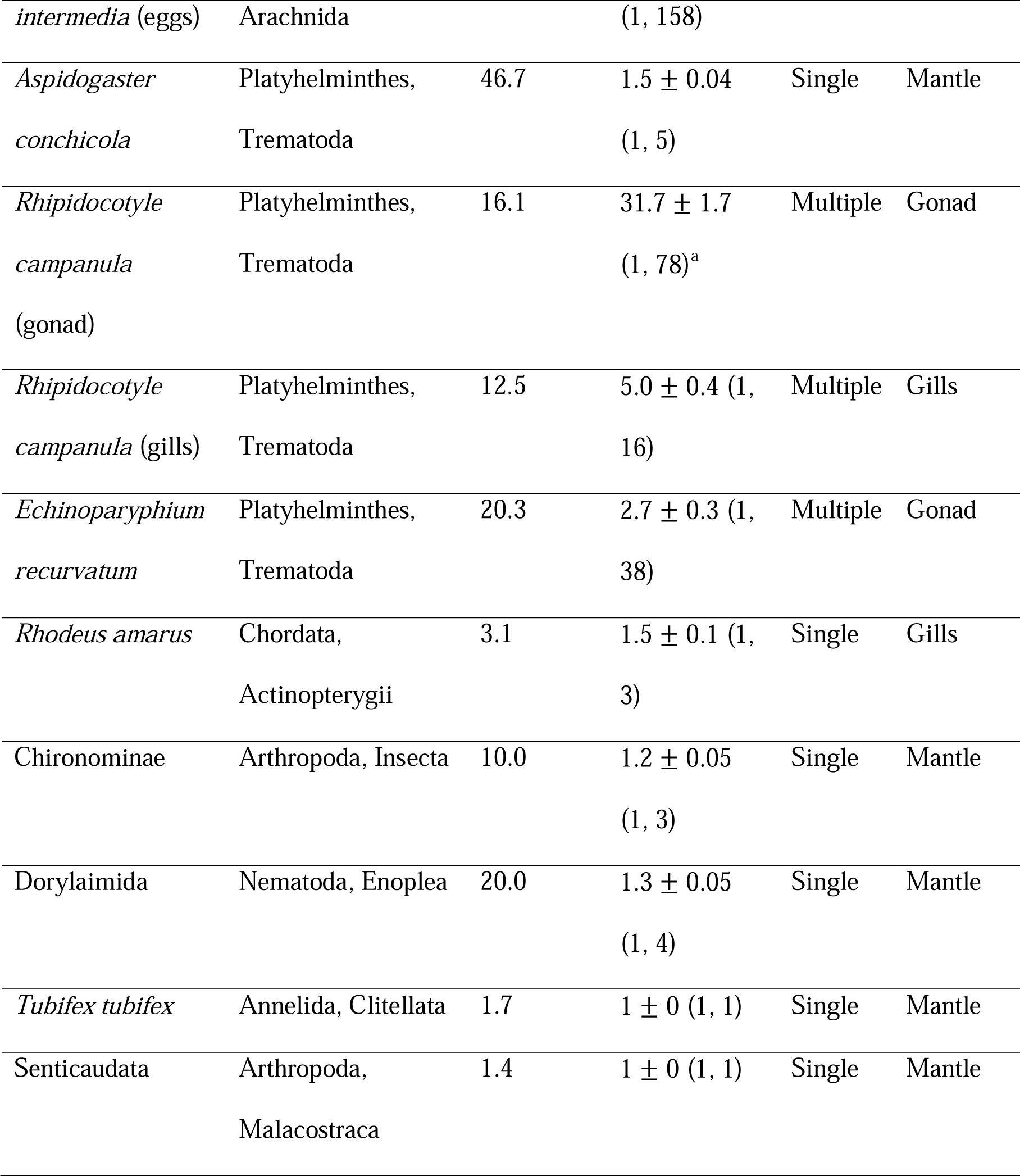
All parasites >1% prevalence identified across 720 *A. anatina*. To capture all possible interactions, species are included as multiple entries if they occur independently in multiple forms or in multiple host tissues (see S1 Supplementary Methods). Mean intensity was calculated only for those mussels with that particular parasite present (Bush et al. 1997). Intensity is a direct count of the number of that parasite in a given host, except for *R. campanula* in the gonad, which was measured as the percentage of the gonad filled with trematode tissue. Life history describes whether the mussel is the only host of the parasite (Single) or whether other hosts are utilised in the parasite life cycle (Multiple), while Location describes the location in the mussel that the parasite is found (mantle, gills or gonad).

Analysis also incorporated five environmental factors as explanatory variables, in two broad categories. The first category (‘mussel quality’) are characteristics of the individual host mussel: length (mm), gravid status (yes/no), and weight. Gravidity refers to the phenomenon of freshwater mussels brooding larval mussels for a period of time in specialist tubes (marsupia) in the outer demibranchs of their gills. A male/female designation was not made, as 94.3% of mussels displayed the female characteristic of marsupia in the outer demibranchs; hence there was likely a significant number of hermaphrodites. Mussel weight was regressed against length given the two variables covaried (R^2^ = 0.91); the residuals from this model (in g) were used as the explanatory variable ‘weight’, to account for weight differences between mussels not explained by length. The second category of environmental variables (‘dispersal influencers’) are those which could affect parasite dispersal to the mussel: invasive zebra mussels (*Dreissena polymorpha*) attached to the host shell (yes/no), and month of the year (12 levels).

### Statistical analysis

Modelling consisted of two complementary methods: Redundancy Analysis and Joint Species Distribution Modelling. Research has shown that multiple approaches should be used to detect community drivers, as methods often have different sensitivities to environmental variables or the nature of the response data (Zhang et al. 2018; Norberg et al. 2019; Ovaskainen et al. 2019). All analyses were carried out using R v.3.6.2 (R Core Team, 2019). Redundancy Analysis was executed using functions available in the package `vegan` (Oksanen et al. 2019), while Joint Species Distribution Modelling was executed with the package `Hmsc` (Tikhonov et al. 2019). For specifics, see S6 Supplementary R Code. For both, modelling utilised the following three matrices. (a) The matrix **Y**_**AB**_ (S3 Supplementary Data 1), which consists of abundance records for all 14 parasite types in 720 sampling units (individual mussels). This was used for the abundance (AB) models. (b) The matrix **Y**_**PA**_, in which **Y**_**AB**_ was modified to only contain zeroes and ones (i.e. records presence-absence only). This was used for the presence-absence (PA) models. (c) The environmental covariate matrix **X** (S4 Supplementary Data 2) which consists of values for the five environmental variables for the corresponding mussels. All procedures outlined below were executed for both the **Y**_**AB**_ and **Y**_**PA**_ matrices.

### Redundancy analysis (RDA)

RDA (Rao 1964) is a form of constrained canonical ordination, and is the traditional method for examining multi-species responses to environmental variables (Zhang et al. 2018; Tikhonov et al. 2019). Prior to analysis, **Y**_**AB**_ was Hellinger-transformed to make the resulting matrix Euclidean (necessary for RDA); of available transformations, this typically performs best (Legendre and Gallagher 2001; Blanchet et al. 2014). No transformation is necessary for **Y**_**PA**_ (Blanchet et al. 2014), though to confirm this an RDA was also run with a Hellinger-transformed presence-absence matrix; there was a <0.2% difference in explanatory power between the two. The results from the untransformed **Y**_**PA**_ are therefore presented.

A standard RDA of **Y**_**AB**_ ∼ **X** (**Y**_**PA**_ **∼ X** respectively) with all default options was performed. Global significance of the model, plus significance of individual canonical axes, were tested for using a permutation test with 999 permutations. To test whether both ‘mussel quality’ and ‘dispersal influencers’ are important correlates of community structure, partial RDAs were executed. The three variables comprising mussel quality were tested for significance, holding the two variables of dispersal influencers constant in the model, and *vice versa*. Finally, forward selection using AIC was executed to determine the specific variables contributing the significant results found from the tests above (see Results). If model AIC values differed by <2, the most parsimonious model was selected (Burnham and Anderson 2004). The explained variation in the global RDA model was then partitioned among those variables found significant by forward selection.

To check for multicollinearity, Variance Inflation Factors were calculated for each explanatory variable in **X** (and each level for categorical variables). All VIFs were <2, confirming an absence of linear dependencies.

### Joint Species Distribution Modelling

Joint Species Distribution Models (JSDMs) were implemented in the Hierarchical Modelling of Species Communities (HMSC) framework (Ovaskainen et al. 2017; Tikhonov et al. 2019); this framework performed the best of any JSDM in extensive simulations (Norberg et al. 2019). This method implements Bayesian probability to predict both individual species’ responses to environmental space, and species-species interactions after accounting for their independent and/or covarying responses to all environmental variables, something beyond the remit of traditional techniques such as C-scores. This is achieved through a latent variable approach, which introduces modellable random variation at the sample level (see Warton et al. 2015; Ovaskainen et al. 2016).

Alongside the **Y**_**AB**_, **Y**_**PA**_ and **X** matrices described above, modelling also incorporated the trait matrix **T** (S5 Supplementary Data 3), describing the values of the two traits for each parasite species (final two columns of Table 1). This matrix can be used to test for the possible influence of these traits on species’ responses to **X**. To allow for the examination of parasite interactions, we modelled mussel host as a random variable with 720 levels.

The model was constructed and run with default `Hmsc` priors (Tikhonov et al. 2019) using 2 MCMC chains, each of 300,000 samples, with the first third of each chain discarded as burn-in and the remainder thinned to every 200^th^ sample. For **Y**_**AB**_, parasites were modelled using a log-normal Poisson distribution, except for *T. tubifex* and Senticaudata, which were modelled using probit regression given they had a maximum intensity of 1. For **Y**_**PA**_, all 14 parasites were modelled using probit regression. The fit and predictive power of the model was assessed through four-fold cross-validation, and model performance was quantified using Tjur’s R^2^ (Tjur 2009), SR^2^ conditional on counts (Tikhonov et al. 2019), and area under the receiver operating characteristic (AUC) (S2 Supplementary Results, Table S2.1).

The variance in the model was partitioned according to the different environmental variables, to examine determinants of community structure. In addition, the variance-covariance matrix **Ω** was calculated to assess potential parasite-parasite interactions (standardised to a correlation matrix, with values between -1 and 1).

## Results

In total, 14 different kinds of parasite were identified (Table 1). All 720 mussels had at least 1 parasite, with a maximum of 10 (4.68 ± 1.61, mean ± s.d.; Fig. S2.1). Parasite prevalence and intensity varied both between parasite type (Table 1), and also throughout the year (Figs. S2.2, S2.3). To explore possible drivers of parasite community structure, two classes of analyses were executed: Redundancy Analysis and Joint Species Distribution Modelling, using both abundance (AB) and presence-absence (PA) models (the input matrices **Y**_**AB**_ and **Y**_**PA**_, respectively).

### Redundancy analysis

The AB model was more successful than the PA model at explaining variation in the parasite matrix (R^2^_Adj_ of 0.22 *vs*. 0.09); however, a global permutation test revealed that environmental covariates played a significant role in influencing the parasite community of a mussel (p = 0.001 in both cases). Further, for both AB and PA models, a test of individual canonical axes revealed that the first four RDA axes were significant, suggesting multiple independent axes of variation in parasite community structure.

All four partial RDAs (testing ‘mussel quality’ factors holding ‘dispersal influencers’ constant, and *vice versa* for AB and PA models) were significant (p = 0.001 in all cases). The partial RDAs indicate that both mussel quality and dispersal influencers provide a valuable contribution to the model, even in the presence of the other group of factors. This was affirmed by forward selection, where environmental covariates from both these groups were selected in the final models. For both AB and PA models, the factors month, length and gravidity were included (Fig. 1).

**Figure 1:**
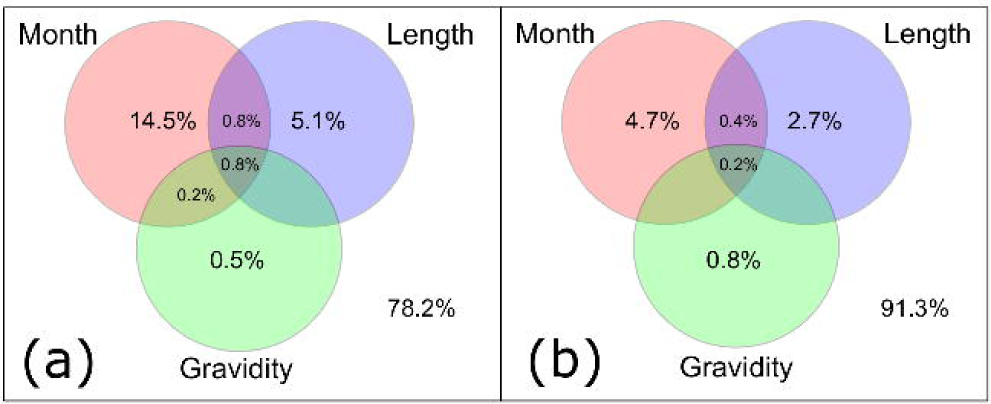
Venn diagrams showing the proportion of variation (as a percentage) explained by the three factors chosen by forward selection, or by a combination of factors. Cells with missing values had an R^2^_Adj_ of < 0. The value not in a circle represents residual unexplained variation. Sizes of circles are not proportional to the amount of variation explained. (a) AB model. (b) PA model.

Fig. 1 shows that in both cases, the factor month explained the greatest proportion of variation, followed by mussel length and then gravid status. However, there were small proportional differences between the two models. Of the variation that is explained, month accounted for 67% in the AB model, but only 54% in the PA model, while length accounted for 23% in the AB model and 31% in the PA model.

### Joint Species Distribution Modelling

Despite the high variation observed in both prevelance and intensity of different parasites between individual mussels and in different months, JSDMs generally performed well when predicting parasite communities (Table S2.1). Analysis of the predicted posterior parameters showed the wide-ranging and variable effects the environmental factors had on the parasites (Fig. 2).

**Figure 2:**
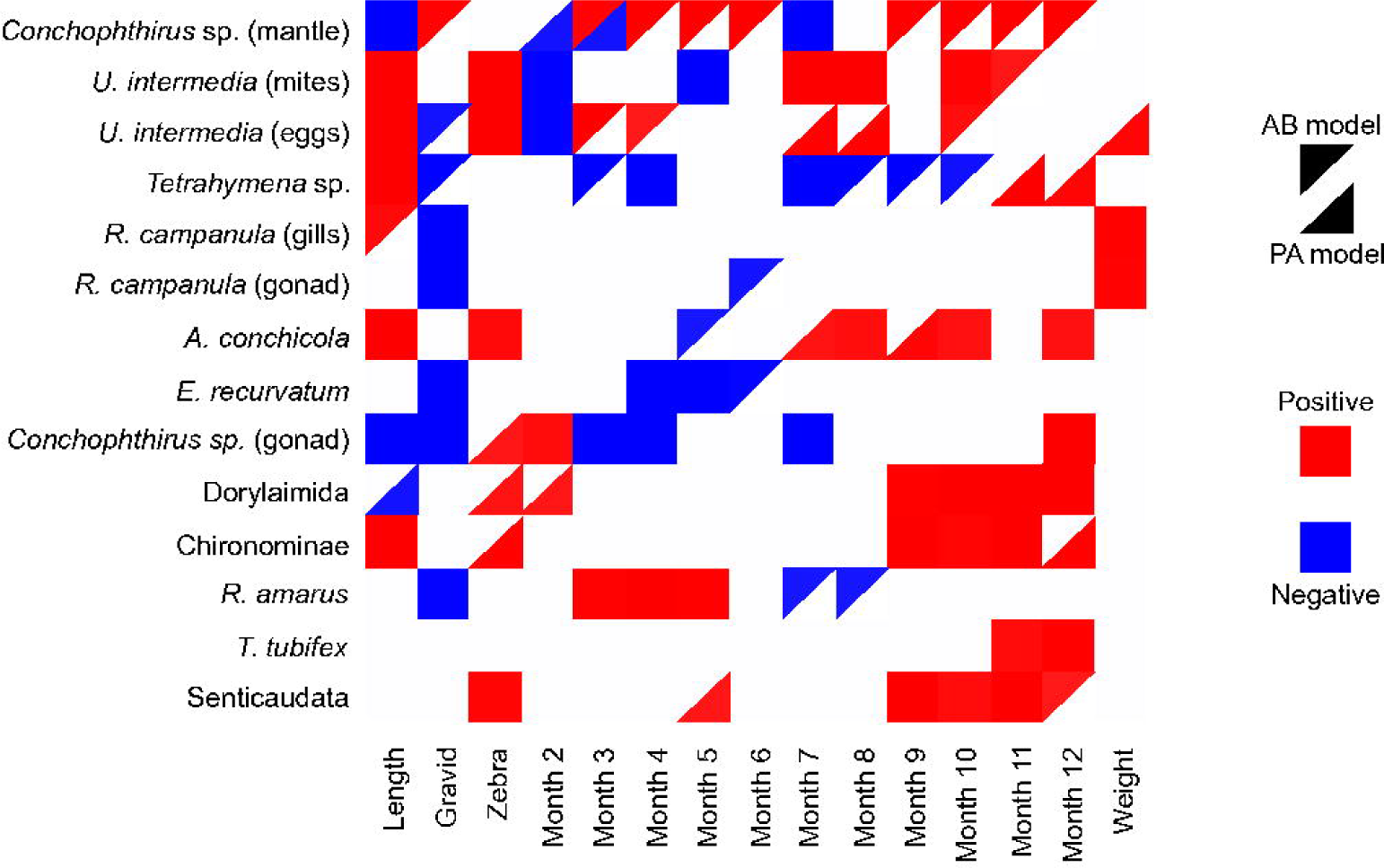
Matrix of β-parameters and their predicted impact on individual parasites, for the abundance (AB) and presence-absence (PA) models. Only effects with > 95% confidence are shown (rest in white). Red indicates positive parasite response to that particular parameter, with blue indicating negative. The upper diagonal of a given square denotes the AB model; the lower denotes the PA model. Note that parameter estimates for categorical factors (Gravid, Zebra and Month) are done with respect to a reference level, so the effect listed here as Gravid is comparing to non-gravid mussels, the factor listed as Zebra is comparing to mussels without zebra mussels, and all Month factors are comparing relative to Month 1. The uninformative Intercept parameter is excluded from the matrix.

Fig. 2 shows that most parasite species were positively or negatively associated with multiple factors of importance. The predictions are highly consistent: with one exception (the square [*Conchophthirus* sp. mantle], [Month 3]), there were no direct contradictions between the AB and PA models. Often, both diagonals of the square are shaded, highlighting the importance of that parameter for both models. The diverse range of effects in the 11 month parameters demonstrates how prevalence and intensity of parasitism varied through the year, often independently for different parasites. Parasites generally responded positively to increased mussel length, and negatively to the mussel being gravid, though there were exceptions. Mussel weight appeared to be a less important factor, but did have a positive impact on the castrating trematode *R. campanula*.

The variation was then partitioned according to the different environmental covariates individually for each parasite (Fig. 3). Averages across all parasites are presented in Table 2.

**Table 2:** The overall percentage contribution of the environmental factors in explaining the variation of the **Y**_**AB**_ and **Y**_**PA**_ matrices.

|  | Month | Length | Gravidity | Zebra | Weight | Random |
| --- | --- | --- | --- | --- | --- | --- |
| AB Model | 50.0% | 11.7% | 15.3% | 3.3% | 3.7% | 16.0% |
| PA Model | 48.3% | 12.1% | 8.6% | 3.9% | 2.4% | 24.7% |

**Figure 3:**
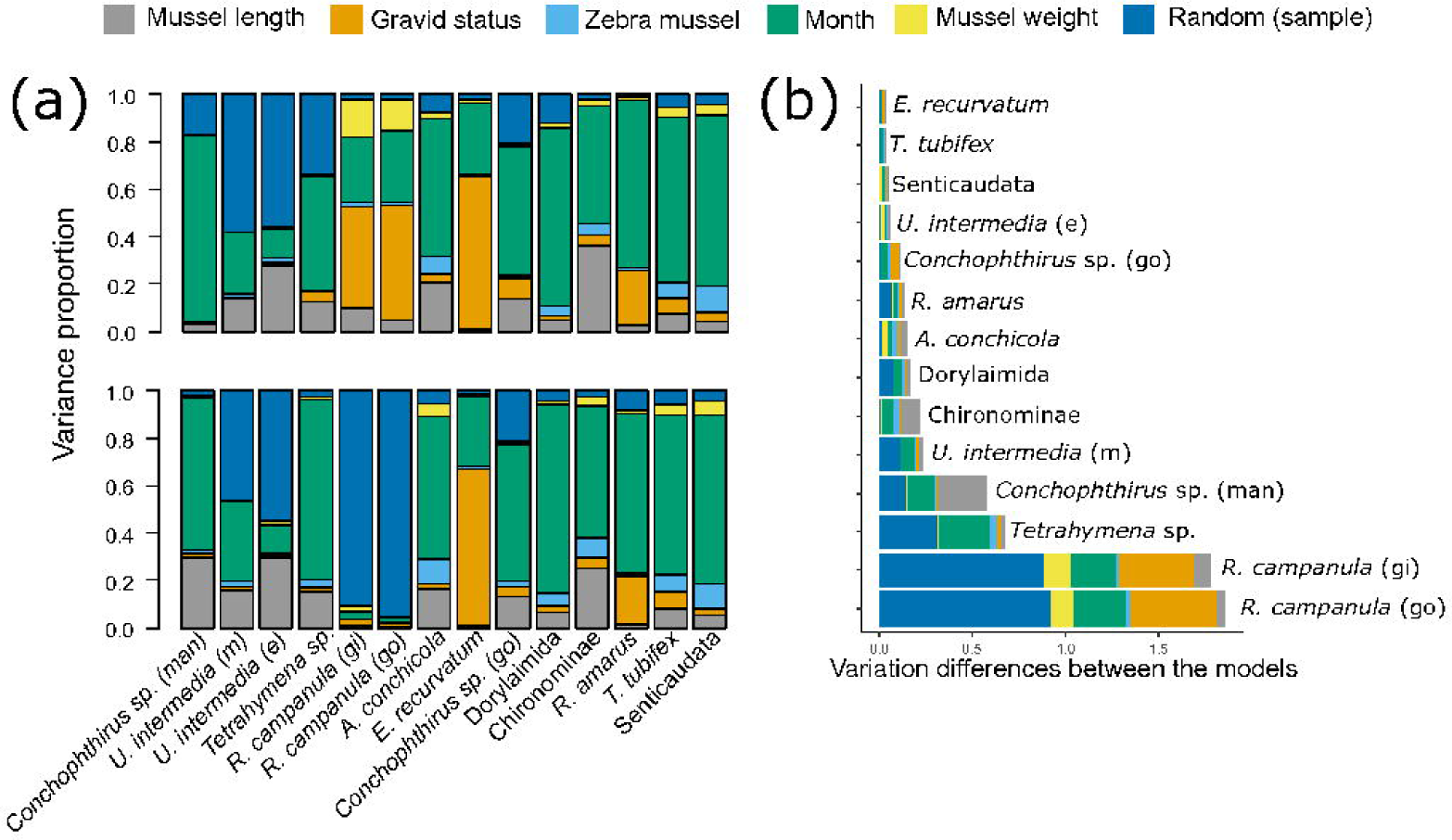
Variance partitioning based on posterior parameter estimates from the JSDM framework. (a) Variance partitioned individually for all 14 parasite species, for the AB model (top) and the PA model (bottom). The colour blue represents the random variation on an individual mussel level not explained by the environmental covariates. (b) Bar plot showing the cumulative differences between the AB and PA models for all 14 parasites. This was calculated as the absolute value of difference for each component of variance, and then visualised in the same colour scheme as (a) and ordered by total difference in variation. This highlights which parasite species had the greatest difference between the AB and PA models (towards the bottom of the chart), and which specific environmental covariates contributed to this difference. e=eggs; gi=gill; go=gonad; man=mantle; m=mites.

Overall, month was the most important environmental factor, accounting for roughly half the variation in the model (Table 2). Length and gravidity also explained a substantial portion of the variation in both models, though overall the PA model had a higher proportion of unexplained variation. However, the importance of these covariates was very different between parasite species (Fig. 3a). Further, there were some marked differences between AB and PA models for individual parasites (Fig. 3b). Particularly, *Conchophthirus* sp. (in the mantle) had more of its variation in abundance explained by month, and more of its variation in presence explained by mussel length. In contrast, the gill ciliate *Tetrahymena* sp. had much more of its variation in presence explained by month, while more variation in abundance was attributed to random variance. Finally, *R. campanula* (both in the gonad and gills) had a vast majority of their variation in presence attributed to random factors; when their abundance was considered, mussel weight and gravid status, in addition to month, became important.

Parasite traits played a role in the differing importance of environmental covariates to different parasites. On average, 42.1% (AB model) and 36.3% (PA model) of the variation between species’ responses was explained by the trait matrix **T**. Particularly, parasite traits were very important in predicting a parasite’s response to whether a mussel was gravid or not (average of 70% of the variation in response to gravidity explained by **T**, Table S2.2).

Finally, the **Ω** matrix of parasite-parasite interactions was inspected, taking into account all other factors in the analysis, and used to construct network diagrams for the AB and PA models (Fig. 4). A liberal criterion of 70% confidence in the interaction was applied prior to inclusion in the matrix, given the small sample size of some parasites. A matrix with a stricter 95% criterion is shown in Figure S2.4.

**Figure 4:**
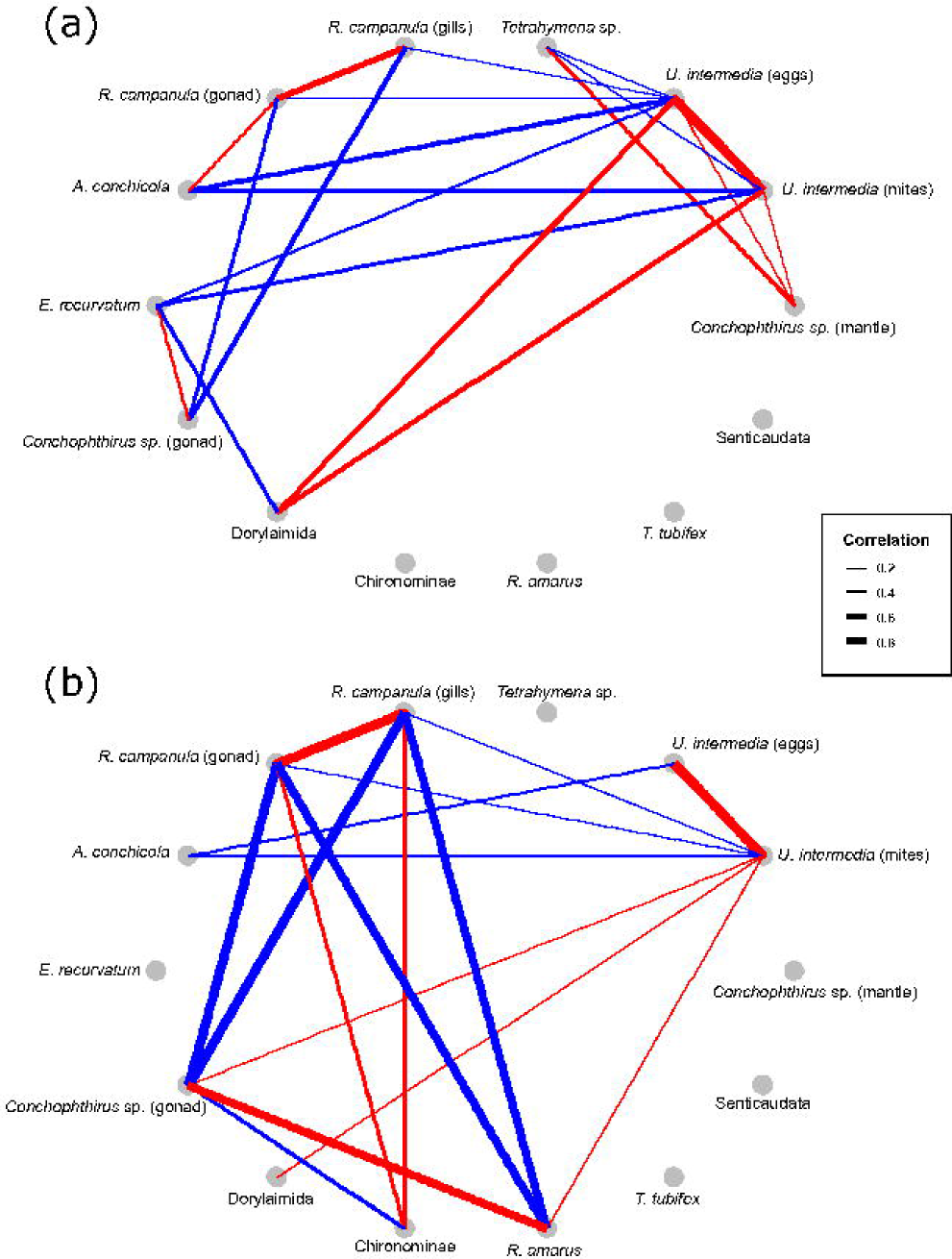
Networks of parasite interactions extracted from the Ω-matrix. Only interactions with > 70% confidence are shown. Red indicates positive correlation, while blue represents negative correlation. Line width is proportional to the strength of the correlation. (a) AB model. (b) PA model.

Fig. 4 shows a diverse range of positive and negative correlations among parasites, even after accounting for all other factors. There were strong positive correlations between the two forms of *U. intermedia* and *R. campanula*, though not for *Conchophthirus* sp. Mites and mite eggs had a large range of negative interactions, both with other parasites in the mantle (*A. conchicola*) and also those in the gonad and gills (*Tetrahymena* sp., *R. campanula, E. recurvatum*). *R. campanula* also had strong negative association with *Conchopthirus* sp. in the gonad, and also with the bitterling *R. amarus* in the PA model. There were also other unexpected potential interactions involving Dorylaimid nematodes, chironomids and *Conchophthirus* sp. in the gonad, especially in the PA model. Additionally, there was less consistency between AB and PA models, with only 20% of interactions having support in both models (in contrast to 59% consistency between the models in their response to the various β-parameters, Fig. 3). Of the 30 interactions in Fig. 4, only 37% of them occurred between parasites occupying the same tissue.

## Discussion

### Predictors of parasite communities in A. anatina

This study modelled the correlation of five environmental covariates (mussel length, weight and gravid status; month and presence of zebra mussels) with the distribution and abundance of parasites and their interactions. The JSDMs had far less random variation present than the RDAs (Figs. 1, 3), likely due to their increased complexity and parameterisation. However, there were very consistent results between the two analytical frameworks. For both AB and PA models, RDA and JSDMs found month, length and gravidity were the three most important factors (Figs. 1, 3; Table 2).

The major driver of parasite community composition in *A. anatina* was month, which suggests propagule pressure is a key determinant of parasite presence and number. As propagule pressure is controlled by both the production of transmission stages (Tavares-Diaz et al. 2014; Rynkiewicz et al. 2019), and how the environment influences the survival of those transmission stages, understanding life-history traits of parasites is a key aspect of predicting infracommunity composition. Because these factors are, in principle, independent of all mussel characteristics, month can be considered an approximately ‘neutral’ factor (*sensu* Gravel et al. 2006). However, a neutral factor does not necessarily imply randomness: our results suggest it may be possible to predict parasite prevalence in a mussel community based on the time of year it was sampled (see also Figs. S2.2, 2.3). Besides parasite population dynamics potentially being predictors of overall prevalence, the consideration of parasite traits is also valuable when considering the parasite infracommunity of a single mussel (Table S2.2). Particularly, parasites occupying the gonad and gills were negatively correlated with mussel gravidity (Fig. 2); considering individual mussel characteristics that influence resource availability could therefore also aid predictions regarding the type of parasites that may infect it.

The importance of mussel-specific resource availability is supported by mussel length and gravidity also being significant predictors of parasite community structure (Fig 1, Table 2). Larger mussels have a larger resource base, and parasites infecting gravid mussels face competition with larval mussels; these may mediate parasite competition, and consequently can be considered ‘niche-based’ factors. Generally, parasites were positively correlated with increased length, and negatively with gravidity in both AB and PA models (Fig. 2), suggesting there is competition for resources both between parasite species, and within a single parasite species. However, positive correlations between parasites and length may also reflect a greater cumulative chance of infection by a parasite, as length increases with age in all but the very oldest mussels (Lundquist et al. 2019). Differentiating between these explanations ultimately requires experimental investigation (Fenton et al. 2014; Poulin 2019). Overall, larger mussels may be at greater risk of parasitism, and gravid mussels may experience reduced parasite pressure (or, conversely, mussels with high parasite loads may be less likely to become gravid). This also highlights the challenge of inferring cause and effect from correlative data, a hurdle typically yet to be overcome in parasite community ecology (but see e.g. Budischak et al. 2016).

The division between ‘niche-structured’ and ‘neutral-structured’ communities is no longer considered a binary separation; communities instead lie on a continuum in which both sets of factors are important (Gravel et al. 2006). However, communities are still commonly discussed as being assembled largely by niche-based or neutral-based factors (e.g. Connolly et al 2014; Mitchell et al 2019). Recent research highlighted that multiple assembly rules can determine the structure of different community modules, both in general communities (e.g. Fournier et al. 2016) and parasite communities (e.g. Williamson et al. 2019). Our research supports this idea that both neutral and niche-based factors are important; further, the relative importance of these factors is parasite-specific, with some parasite species being driven more by niche-based factors (abundances of *R. campanula*, and *E. recurvatum* in both models), and others being driven more by neutral factors (Fig. 3). Consequently, it is erroneous to apply a broad categorisation regarding what structures a *community*, as it may be composed of different species all being influenced independently by a range of factors to varying degrees.

Overall, our results demonstrate that predicting parasite community structure requires an integrative approach, incorporating parasite and host traits alongside the wider environmental and temporal context (Hellard et al. 2015). Understanding such nuances provides a new and exciting challenge in parasite community ecology.

Our results suggest that within-host parasite interactions may also have an important structuring role, though the observed associations (Fig. 4) may be due to parasites responding in similar or opposite ways to an unmeasured environmental covariate (Blanchet et al. 2020). The observed interactions are also striking for the divergences between the AB and PA models, suggesting that in some cases, parasites were limiting (or facilitating) the *numbers* of other parasites, while in others they were limiting or facilitating the *presence* of other parasites. For example, the embryos of bitterling fish (*R. amarus)* had a negative relationship with *R. campanula* in the PA model (Fig. 4b). Given female ovipositing bitterling have previously shown sensitivity to host quality (e.g. Smith et al. 2001; Mills and Reynolds 2002), they may detect lowered host quality due to trematode presence and choose instead to oviposit in uninfected mussels. Conversely, *U. intermedia* had negative relationships with both gill ciliates and trematodes in the AB model (Fig. 4a), suggesting that while presence is not affected by this interaction, both could directly or indirectly compete for resources, thus limiting each other’s abundance.

There were also a high number of interactions between parasites in different tissues (Fig. 4), in contrast to patterns in other studies (e.g. Dallas et al. 2019). However, this fits with previously described theory. An interactive parasite community is more likely to be generated by a high prevalence of each parasite than by a site with high parasite richness overall but low richness in a single host (Dove 1999). Previous studies include multiple hosts in their models, which may not be competent for all parasites. This reduces infracommunity richness relative to total parasite richness, and may increase overall parasite variation to the extent that interactions within a single host species cannot be detected. Conversely, our study utilised only one host species, and thus by default the entire parasite community is guaranteed to be competent in a given individual. Our focus on a single host species suggests that parasites with different life-history strategies may interact across multiple tissues more than previously reported (but see Ferrari et al. 2009; Eidelman et al. 2019).

### Presence-absence versus abundance data: implications for modelling

Overall predictions were similar between PA and AB models (Fig 1, Table 2). However, predictions for several parasite species individually had highly divergent outcomes between the two model sets (Fig. 3b). There were also broad visual differences between the AB and PA parasite interaction networks (Fig. 4). Importantly, there were no explicit contradictions between the two models (e.g. where one model finds a factor to be positively associated with a parasite, and another finds it to be negative), suggesting that model differences are being driven by varying sensitivity to different characteristics of the data, rather than necessarily displaying ‘incorrect’ information. This may still have severely negative consequences for predicting the impacts of parasitism on host populations. For example, the greatest difference between AB and PA models was observed for *R. campanula* in the gills and gonad (Fig. 3b), a castrating trematode that rapidly fills the host with asexually-produced sporocysts and cercariae (see S1 Supplementary Methods). Inspecting the PA model alone, *R. campanula* had little explanation for variation in its presence, leading to the conclusion that it has little relationship with the measured mussel characteristics. However, the AB model revealed a significant negative correlation with mussel gravidity (Figs 2, 3a), likely reflective of the fact that a higher intensity of infection can lead to castration of the host mussel (Taskinen et al. 1997). If only presence-absence data was available, as it common in parasite community studies, this interaction might be overlooked, though it can have severely negative population-level consequences for the host mussel (Taskinen and Valtonen 1995). This study therefore highlights the importance of using abundance data to accurately characterise host-parasite interactions.

Previous research has explored whether presence-absence data is also useful in predicting abundances in ecological communities, and is relevant here to explain the divergences of AB and PA models. Joseph et al. (2006) showed that abundance data generally outperformed presence-absence data in assessing population declines and reductions in area of occupancy. However, for rare or cryptic species with low abundance or detectability, presence-absence measures performed as well as (and sometimes better than) abundance. Podani et al. (2013) showed that presence-absence measures are useful when considering species composition, but (intuitively) less so when assessing more subtle quantitative trends. These results suggest presence-absence data may be appropriate for modelling species with low abundance (where counts do not diverge far from binary presence/absence anyway), while species with higher abundance require explicit abundance measures. Indeed, in our study, the five parasites with the greatest divergence between AB and PA models (Fig. 3b) are five of the six parasites with the highest intensities in mussels (Table 1). Further, those species with high intensities (e.g. *Conchophthirus* sp., *U. intermedia, Tetrahymena* sp.) had many more potential interactions in the AB model that were absent in the PA model (Fig. 4a), while lower-intensity parasites (*R. amarus*, Chironominae) had interactions in the PA model but not the AB model (Fig. 4b). Abundance data may therefore reflect habitat *quality* (i.e., how much of a particular parasite can a mussel support?), while presence-absence data reflects habitat *suitability* (i.e. can a mussel support that parasite?) (Gutiérrez et al. 2013).

Presence-absence data may still be useful for detecting broad prevalence trends and species’ interactions, though recent theoretical work has demonstrated multiple issues with it (Blanchet et al. 2020). Sometimes, the same factors may govern both distribution and abundance, making presence-absence data a useful and simple way of assessing community trends (e.g. Gutiérrez et al. 2013). However, our work suggests that, for species with high abundance, presence-absence data may be highly misleading and lead to incorrect and potentially damaging ecological inference. Parasite species with high prevalence and variable levels of intensity within a host population may have contrasting outcomes at both an individual host and population level (Hurd 2001; Lafferty and Kuris 2009); consequently, understanding the factors driving their assembly in individual hosts is vital to predicting parasite effects. We strongly recommend that, where possible, abundance data is incorporated into studies of parasite community ecology, as it has the capacity to explain greater levels of variation in multiple analytical frameworks (e.g. Fig. 1, Table 2). An increased emphasis on invertebrates may facilitate this, especially with the advent of new methods to accurately quantify infection in these hosts (e.g. Brian and Aldridge 2020). This paper has demonstrated the correspondence of parasite communities with both niche- and neutral-based factors, alongside a range of possible parasite interactions, all from a single host species at a single site. Given the complexity of this picture, which leaves aside multiple sites or host species, we show that data needs to be as nuanced and as fine-scale as possible to expand our understanding of parasite community ecology.

## Supporting information

S2 Supplementary Results

## Acknowledgements

JIB is supported by a Woolf Fisher Scholarship. DCA is supported by a Dawson Lectureship from St Catharine’s College, Cambridge. We thank Camilla Campanati, Sebastian Dunne and Christine Ellis for assistance with sampling, and Miriam Shovel for laboratory and editorial assistance. The authors declare no conflict of interest.

