## Supplementary material for "Abundance data applied to a novel model invertebrate host sheds new light on parasite community assembly in nature": S1 Supplementary Methods

*Mussel size through sampling period*

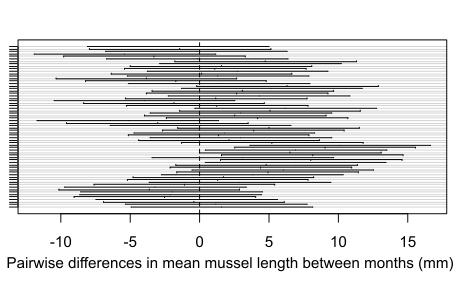

Figure S1.1: Pairwise differences (y-axis) between the mean mussel lengths for all 12 sampling months, beginning with all pairwise comparisons for the first month (top of y-axis) and moving progressively down. Black bars indicate a 95% C.I. for the difference between two months; note that 72 of the 78 C.I. overlap 0.

There was no evidence that mussel size changed significantly throughout the year, or that mean size was affected by our sampling. While there were differences between months (F_(11, 108)_ = 4.17, p < 0.0001), this was driven only by slightly smaller mussels in months 5 and 6 (Tukey post-hoc test, p < 0.05 for 6 of the 21 pairwise comparisons for these two months), consistent with timings of recruitment of new *A. anatina* into the population. Fig. S1.1 shows no clear pattern of sampling causing a consistent reduction in overall mussel size.

*Mussel dissection and parasite isolation*

Following collection, mussels were transported back to the laboratory in 10 L buckets in river water. In the laboratory, mussels were held at 8 °C under aeration, for a maximum of 72 h before dissection. Immediately prior to dissection, mussels were rinsed under cold fresh water while holding the valves gently shut to remove any organisms on the shells, and had their maximum length recorded to the nearest 0.5 mm with Vernier callipers.

All tissues of the mussel were inspected in systematic fashion. In all stages of dissection, both the presence and the number (i.e. abundance) of all parasites found were recorded, and subsequently identified to the finest possible taxonomic resolution.

Mussels were cut open by gently inserting a scalpel between the valves and slicing the posterior and anterior adductor mussels over a transparent petri dish, to allow all fluids from the mantle cavity to be collected. These fluids were then inspected under a GXMMZS0745-T stereomicroscope at 16$\times$ magnification. Following this, the mussel was placed under the stereomicroscope and the mantle and labial palps on both sides were systematically searched. The inner and outer demibranchs (gills) were then carefully removed individually, and the filaments gently teased apart under 16$\times$ magnification to record all the parasites present in the gills. The visceral mass was gently removed by cutting the connective tissue at each end, and was briefly placed aside, to expose the pericardial cavity, which was also dissected under 16$\times$ magnification. The visceral mass was then cut open with a scalpel at the posterior end where the gonads were located, and examined: samples of gonad tissue were removed with tweezers and pressed between two microscope slides to create a squash ~10 mm in diameter. These samples were inspected under a GXML3200 compound microscope at 40$\times$ magnification. In addition, each sample had three photos taken with a GXM HiChrome-S camera attached to the microscope to quantify infection with digenean trematodes.

The dissecting tray and all equipment were then washed before proceeding to the next mussel. Mussel shells were dried fully and weighed to the nearest 0.001 g, and mussel tissue was dried to constant mass (nearest 0.001 g).

Following all dissections, the water remaining in the transport and holding buckets was stirred thoroughly to agitate any sediment, and three 10 mL samples were taken from the bottom of each bucket and inspected under 16$\times$ magnification, to confirm that storage in the buckets did not induce parasites to leave the mussels and affect results. These samples contained large gammarid amphipods (which were commonly observed on the exterior of mussel shells) and very rarely oligochaetes and leeches, which were never observed inside mussels and were also likely present via exterior attachment. No mussel parasites were observed in these samples, which suggests the communities observed upon dissection were consistent with those that were present at collection.

*Parasite identification*

Broadly, the major parasite groups observed were ciliates, mites, aspidogastrean trematodes, bucephalid trematodes, echinostomatid trematodes, bitterling embryos, nematodes, chironomid larvae, oligochaetes, and amphipods. Identifications were made to the finest taxonomic resolution possible for each of these groups. As additional verification, all previous parasite records for this and closely-related mussel hosts were inspected (summarised in Brian and Aldridge 2019 [Table S3]), and significant deviations from previous records carefully checked.

Ciliates were isolated from the tissues they appeared in, stained with methyl-green pyronin (to highlight the nuclear material), then mounted and inspected under 400$\times$ magnification following Foissner and Berger (1996). Two species of ciliates were clearly distinguished. The first was observed in the mantle (localised particularly on the labial palps), and was identified as *Conchophthirus* sp. (Fig. S1.2). The specific species is uncertain, as there are few explicit taxonomic keys available for unionid ciliates. One of the most useful is that of Kidder (1934); the mantle species observed here matches the life history strategy (with particular note of the location on the labial palps) of *C. anodontae*, but the location of the macronucleus and distribution of endoplasmic granules align very closely with that of *C. curtus*. Given this uncertainty, this mantle ciliate has been identified to genus only.

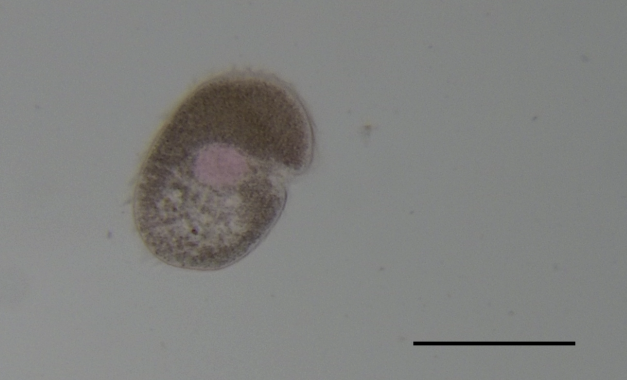

Figure S1.2: *Conchophthirus* sp. Note the macronucleus (pink). Scale bar 100 μM.

The second ciliate species was observed in the gills, and identified as *Tetrahymena* sp., possessing all the broad characteristics of this genus (Fig. S1.3). The macronucleus of this species proved difficult to stain, with no clear macronuclear region. However, there were small pink deposits through the cytoplasm, possibly representing the nuclei of ingested host cells (see Lynn et al. 2018). There appear to be no keys for the identification of *Tetrahymena* species, and so the classification was only done to genus level. Given that, to our knowledge, these are first ciliate records from *A. anatina* (see Brian and Aldridge 2019), it is possible and even likely that one or both are novel species.

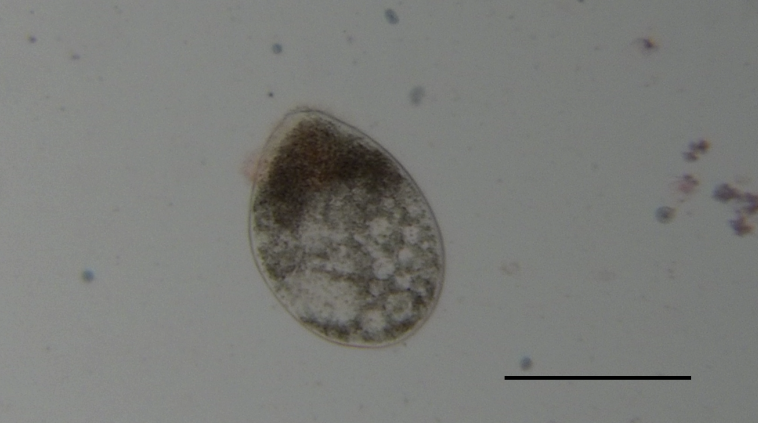

Figure S1.3: *Tetrahymena* sp. Note the absence of clear staining. Scale bar 150 μM.

The gonadal pore and leg segmentation of mites were inspected under 400$\times$ magnification and compared with the keys of Vidrine (1986), in addition to the ecological information of Baker (1988). Mite eggs were also observed; these were not identified directly, but inferred to be of the same species given the congruence of a linear pattern of occurrence on the mantle for both larval mites and mite eggs. Mites were identified as *Unionicola intermedia* (Fig. S1.4). The mites present were typically larval, as the adults are transient and do not spend extended periods of time in the mussel, but adults and later nymphal stages of the mites were also observed. Mites and mite eggs were typically on the mantle, though one nymphal stage was also occasionally observed on the gill margins.

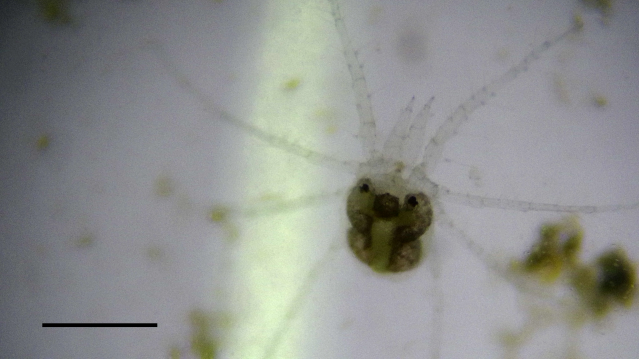

Figure S1.4: *Unionicola intermedia* adult, from mantle cavity of *A. anatina*. Scale bar 750 μM.

Identification of: (a) aspidogastrean trematodes; (b) bucephalid trematodes; and (c) echinostomatid trematodes, was through consultation with (a) Huehner and Etges (1977), Schell (1985), Alves et al. (2015); (b) Taskinen et al. (1991), Gibson et al. (1992); and (c) Conn and Conn (1995), Molloy et al. (1997), Sohn (1998); respectively. Aspidogastrean trematodes were identified as *Aspidogaster conchicola* (Fig. S1.5). These have a simple life history with a single host (the mussel). Both juvenile and adult *A. conchicola* were observed in the mantle cavity under the visceral mass. Bucephalid trematodes were identified as *Rhipidocotyle campanula* (Fig. S1.6). These have a complex life cycle with three hosts; they utilise mussels as their first intermediate hosts (see Taskinen et al. 1997 for a summary of the life cycle), with sporocysts and cercariae present in the mussel gonad. These fill the gonad, and can induce complete castration in their mussel hosts. Cercariae are then released to infect the next intermediate host in the life cycle. Echinostomatid trematodes were identified as *Echinoparyphium recurvatum* (Fig. S1.7). These trematodes also have a complex life cycle with three hosts (see Molloy et al. 1997), and utilise the mussel as a second intermediate host. Metacercariae of this species were observed in the mussel gonad.

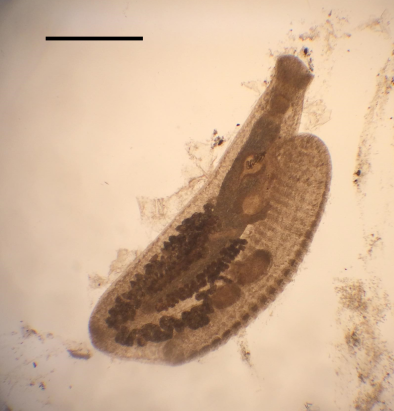

Figure S1.5: Juvenile *Aspidogaster conchicola*. Scale bar 250 μM.

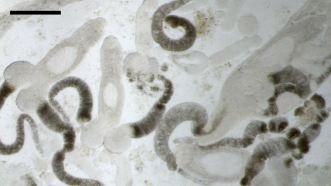

Figure S1.6: *Rhipidocotyle campanula* cercariae. Scale bar 250 μM.

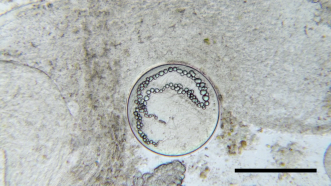

Figure S1.7: *Echinoparyphium recurvatum* metacercaria. Scale bar 150 μM.

Bitterling are freshwater fish that lay their eggs in the gills of freshwater mussels, where they constitute a stress both in terms of physical disfiguration and oxygen competition (Reichard et al. 2006; Methling et al. 2018). The bitterling embryos were readily identified as *Rhodeus amarus*, given this is the only sympatric fish with this life history strategy (Reynolds et al. 1997; Aldridge 1999).

Chironomid larvae and nematodes were identified as far as possible by isolating them from the mussel mantle and inspecting their head structures under 40$\times$ (chironimids) or 400$\times$ (nematodes) magnification, respectively. In both cases, it is difficult to get fine taxonomic resolution, particularly for nematodes as it is likely that the nematodes observed inside mussels are larval (McElwain et al. 2019), and are therefore missing many of the diagnostic characteristics of adults. Literature consulted included Stewart and Loch (1973) and Pinder (1986) for chironomids, and Abebe et al. (2006), Gagarin and Gusakov (2016) and McElwain et al. (2019) for nematodes. Chironomids were classed as subfamily Chironominae, while nematodes were classed as order Dorylaimida (Fig. S1.8); it should be noted that in both of these cases, it is possible that there were multiple species within these groupings.

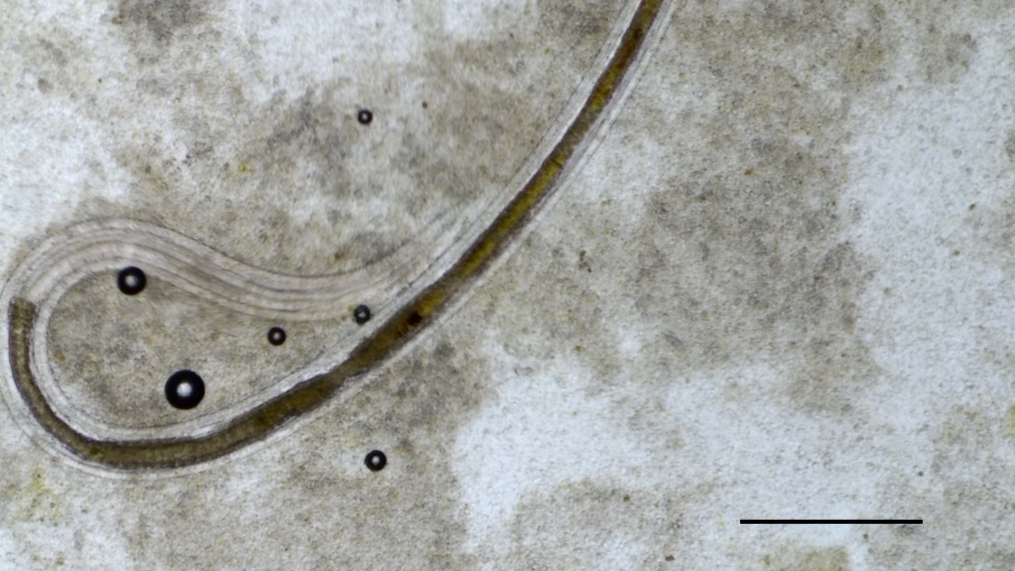

Figure S1.8: Head and start of body of a Dorylaimid nematode. Scale bar 300 μM.

All oligochaetes had the same appearance, which did not match the appearance of any previously recorded oligochaetes in the mantle of freshwater mussels (see Table S3 of Brian and Aldridge 2019). However, the oligochaetes seen were strikingly similar to *Tubifex tubifex* (compared through online taxonomic resources such as the Marine Life Information Network [marlin.ac.uk] and the World Register of Marine Species [www.marinespecies.org], both of which include freshwater species), and distribution patterns also match *T. tubifex*, and so the oligochaetes observed have been assigned this taxonomic classification.

Gammarid amphipods were also occasionally observed in the mantle. Given the freshwater environment in which these amphipods occurred, they have been assigned to the suborder Senticaudata (Lowry and Myers 2013), but further taxonomic classification was not attempted.

In addition to these parasites, several others were observed at frequencies <1%, including single observations of different leech species as well as echinostomatid and bucephalid trematodes. These were not included in the final parasite matrix, as models become increasingly difficult to fit with excess zeros (Ovaskainen et al. 2016; Lammel et al. 2018), and their extreme rarity suggests low importance (and predictive power) in mussel parasite communities.

*Construction of the parasite matrix*

In three cases, the same parasite species was observed in multiple locations in the mussel, or in different life-history stages. First, *Unionicola intermedia* was present as both mites and eggs in the mantle. Second, live *Conchophthirus* sp. was observed both in the mantle but also occasionally in the gonad. It is unclear how these ciliates ended up in the gonad; given their localisation on the labial palps, which coveys food to the mussel’s mouth, they were perhaps accidentally transported into the visceral mass. Third, *Rhipidocotyle campanula* occurred both as sporocysts and cercariae in the gonad, but also as sporocysts running transversely through the gills. In all three of these cases, observations of different forms of the same parasite appear to be independent (for example, cases were observed where there was *R. campanula* present in the gonad but not gills, cases where it was in the gills but not the gonad, cases where both were observed, and cases where neither were observed).

As a result, we believe it is an over-simplification to conglomerate presence or abundance recordings into one category. For example, bitterling (who lay their embryos in the gills) may react negatively to *R. campanula* sporocysts in the gills, but sporocysts in the gonad may have no effect. Given the purpose of this study is to determine possible drivers of parasite community structure, important parasite-parasite interactions could be missed by failing to include life-history forms or different locations of the same parasite. The importance of not grouping potentially independent parasite types has been recently emphasised (Poulin 2019).

Therefore, the parasite matrix (**Y_AB_**) was constructed with 720 rows (sampling units, the mussels) and 14 columns, as follows: *Conchophthirus* sp. (mantle); *Conchophthirus* sp. (gonad); *Tetrahymena* sp.; *Unionicola intermedia* (mites); *Unionicola intermedia* (eggs); *Aspidogaster conchicola*; *Rhipodicotyle campanula* (gonad); *Rhipidocotyle campanula* (gills); *Echinoparyphium recurvatum*; *Rhodeus sericeus*; Chironominae; Dorylaimida; *Tubifex tubifex*; Senticaudata. See Table 1 (main text) for a summary of occurrences.
