## Supplementary material for "Abundance data applied to a novel model invertebrate host sheds new light on parasite community assembly in nature": S2 Supplementary Results

A) **Supplementary Figures**

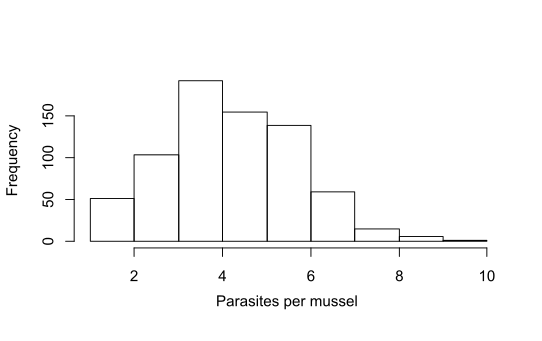

Figure S2.1: Histogram showing the distribution of parasite frequency counts per mussel.

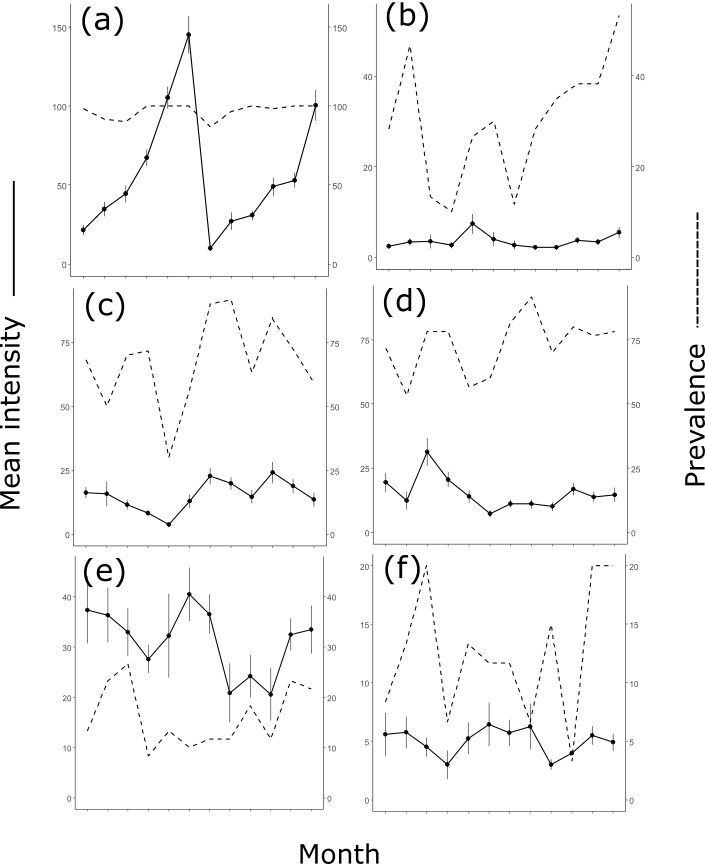

Figure S2.2: Mean intensity ($\pm$ S.E.) and prevalence for: (a) *Conchophthirus* sp. (mantle); (b) *Conchophthirus* sp. (gonad); (c) *U. intermedia* (mites); (d) *U. intermedia* (eggs); (e) *R. campanula* (gonad); and (f) *R. campanula* (gills). Note that each y-axis possesses a different scale.

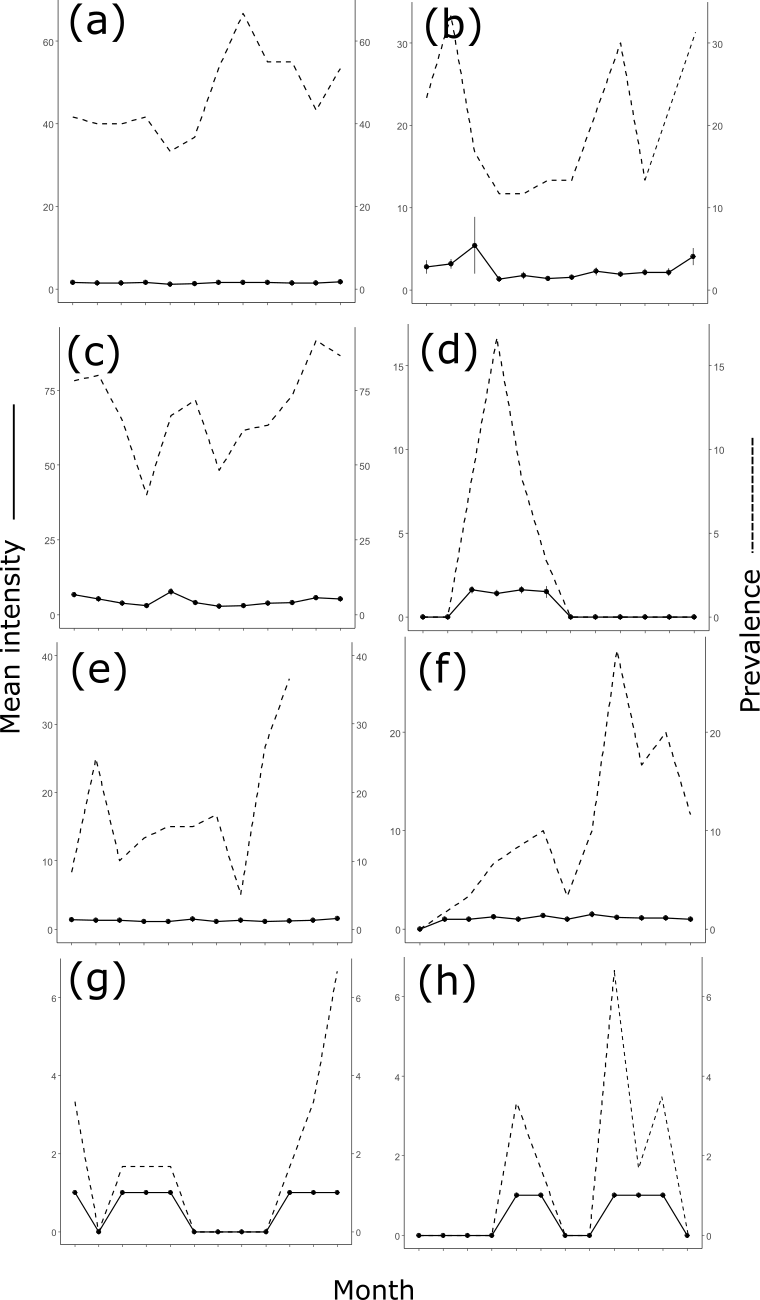

Figure S2.3: Mean intensity ($\pm$ S.E.) and prevalence for: (a) *A. conchicola*; (b) *E. recurvatum*; (c) *Tetrahymena* sp.; (d) *R. sericeus*; (e) Dorylaimida; (f) Chironominae; (g) *T. tubifex*; and (h) Senticaudata. Note that each y-axis possesses a different scale.

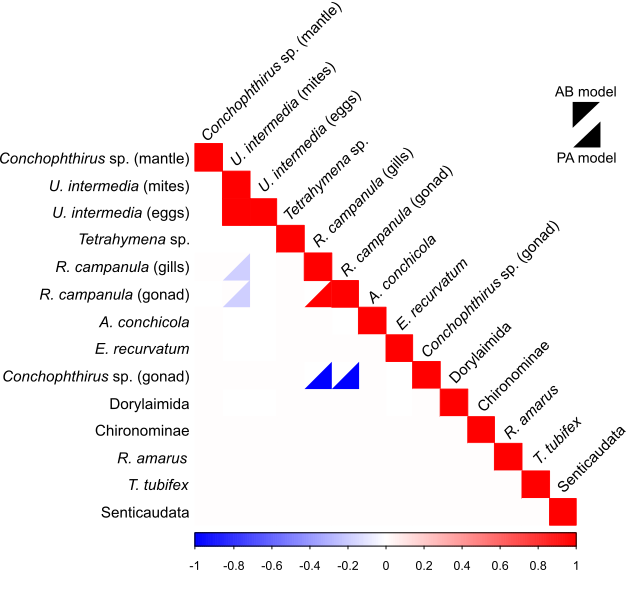

Figure S2.4: Ω-matrix of parasite-parasite interactions. Only interactions with > 95% confidence are shown in the matrix. The blue-red scale represents the correlation coefficient, where 1 (dark red) indicates perfect positive correlation, and -1 (dark blue) indicates perfect negative correlation. The upper diagonal of a given square denotes the AB model, while the lower denotes the PA model.

B) **Supplementary Tables**

Table S2.1: Assessment of predictive power for fitted JSDMs (mean $\pm$ σ in each case)

| **Model** | **AUC** | **Tjur’s R^2^** | **SR^2^** |
| --- | --- | --- | --- |
| Abundance | 0.697 $\pm$ 0.066 | 0.062 $\pm$ 0.038 | 0.146 $\pm$ 0.135 |
| Presence-absence | 0.681 $\pm$ 0.075 | 0.067 $\pm$ 0.052 | N/A |

Table S2.1 shows generally good performance of the fitted models at predicting the data, as they perform significantly better than chance at predicting presences and absences (equivalent to an AUC of 0.5) and have a clear difference between the mean fitted values for presences and absences (Tjur’s R^2^). In addition, the abundance model performed reasonably well at ranking abundances (SR^2^). In all of these cases, more common parasites had better measures of predictive power, and averages were brought down by more incidental parasites, though in all cases values were still > 0 (Tjur’s R^2^, SR^2^) or > 0.5 (AUC).

Table S2.2: Exploring how much of the variation in parasite species’ responses to individual environmental covariates can be explained by the trait matrix **T** (life history of the parasite, and location of the parasite in the mussel).

| **Environmental Covariate** | **AB model** | **PA model** |
| --- | --- | --- |
| Length | 47.2% | 38.9% |
| Gravid Status | 75.4% | 64.1% |
| Zebra mussel presence | 29.9% | 20.4% |
| Month | 41.4% | 36.6% |
| Weight | 16.5% | 21.7% |
| Average | 42.1% | 36.3% |
